# Synergistic effect: a common theme in mixed-species litter decomposition

**DOI:** 10.1101/807537

**Authors:** Jun Liu, Xiaoyu Liu, Qingni Song, Fenggang Luan, Hui Wang, Yalin Hu, Qingpei Yang

**Affiliations:** Jiangxi Provincial Key Laboratory for Bamboo Germplasm Resources and Utilization, Jiangxi Agricultural University, Nanchang 330045, China; Stable Isotope Center, College of Forestry, Fujian Agriculture and Forestry University, Fuzhou 350002, China; 2011 Collaborative Innovation Center of Jiangxi Typical Trees Cultivation and Utilization, Jiangxi Agricultural University, Nanchang 330045, China; College of Resources and Environment, Shandong Agricultural University, Taian 271018, China

**Keywords:** Non-additive effect, litter mixture, litter quality, synergistic effect, litter decomposition, meta-analysis

## Abstract

- Litter decomposition plays a key role in ecosystem nutrients cycling, yet, to date science is lacking a comprehensive understanding of the non-additive effect in mixing litter decomposition.
- In order to fill that gap, we compiled 69 individual studies for the purpose of performing two sub-meta-analyses on the non-additive effect.
- Our results show that a significantly synergistic effect occurs at global scale with the average increase by 2-4% in litter mixture decomposition; In particular, low-quality litter in mixture shows a significantly synergistic effect, while no significant change is observed with high-quality species. Additionally, the synergistic effect turns into the antagonistic effect when soil fauna is absent or litter decomposition enters into humus-near stage. In contrast to temperate and tropical areas, studies in frigid area also show a significantly antagonistic effect.
- Our meta-analysis provides a systematic evaluation of the non-additive effect in decomposition mixed litters, which is critical for understanding and improving the carbon forecasts and nutrient dynamics.

## Introduction

Litter decomposition is a central component of biogeochemical cycles in ecosystems. The rate of decomposition controls nutrients returns and energy flow, which regulates atmospheric carbon emissions, soil organic matter composition, and supply of based mineral nutrients to flora (Schneider *et al*. 2012). Thus, litter decomposition influences ecosystem primary productivity (Bradford *et al.*, 2016). Over past few decades, many studies about litter decomposition have focused on single litter decay (Gartner & Cardon, 2004), which resulted in a profound exploration of the connection between decomposition rate and its impact factors. However, in many ecosystems, litter is generally a mix of multiple species. Previous studies suggest that those various litter species interact with each other during their decomposition (Ball et al. 2008; Gessner *et al.*, 2010), implying that the decomposition rate of litter mixture is different from that of single litter. To deeply discern how the litter mixture impacting the decomposition is essential to understand the carbon and nutrients cycles.

When different litter species mix together, their decomposition rates generally do not equal to the arithmetic mean value, i.e., the expected decay rate, between single litter species (Fig. S1) (Gartner & Cardon, 2004; Steinwandter *et al.*, 2019). Instead, one of two potential results usually develop: synergistic effect and antagonistic effect (Collectively referred to as non-additive effect) (Fig. S1). In general, the chemical characteristics of litter, decomposer activity, as well as other environmental factors often constrain decomposition rate (Berg. B, 2014). In litter mixture, the underlying mechanisms of synergistic effect are mainly from three aspects: the nutrients (such as nitrogen) transfer from high-quality litter species to low-quality litter, the complementary effects of soil fauna and decomposers, and the improvement of microclimatic conditions (Schimel & Hattenschwiler, 2007; Tiunov, 2009; Madritch and Cardinale 2007). Conversely, the antagonistic effect is often induced by the enhancement of microbial nutrient immobilization for litter with poor nutrients or the inhibitory secondary compounds released from low-quality litter species (Hättenschwiler et al., 2005; Montané 2013).

Although experiments concerning the mixture effects on litter decomposition rates have been conducted in numerous individual studies, the conflicting results have hampered any possibility to draw general conclusions (Li *et al.*, 2016). Most of conflicts could mainly result from the significant differences of influencing factors related to litter decompositions in experimental design (Barbe et al. 2018, Leroy et al. 2018, Zhao et al. 2019). Thus far, those various studies on mixed decomposition were all performed in different ecosystems across different climate regions with decomposition durations varying from several weeks to even as long as several years. Moreover, the various mesh sizes used in litterbag and microcosm, two methods that the most commonly employed in litter decomposition, often make the results hard to compare. As such, for the past several years, the scientific community has been requesting a meta-analysis in order to determine a general global pattern (Gartner & Cardon, 2004; Gessner *et al.* 2010).

While studies throughout the literature have summarized and analyzed the non-additive effect in mixed litter decomposition (Gartner & Cardon, 2004; Li *et al.*, 2016), there still remains a lot of unanswered questions. In order to better understand how the mixing influences the decomposition rate of litter, we employed a meta-analysis that built upon the system analysis done by Gartner and Cardon (2004). Finally, 70 individual studies were compiled to perform a global analysis (Note S1). If the litter mixing effects occur, we also aimed to explore seven detailed questions following:

1. How does the climate influence the effect of mixing on decomposition?
2. How does the ecosystem influence the effect of mixing on decomposition?
3. How does the mesh size of litterbag influence the effect of mixing on decomposition?
4. How does the period of experiment influence the effect of mixing on decomposition?
5. How do the evenness and richness of species influence the effect of litter mixing on decomposition?
6. How does lignin content (AKA litter quality) influence the effect of mixing on decomposition?
7. How do the leaf type (broad/needle) and habit (evergreen/deciduous) influence the effect of mixing on decomposition?

## Materials and Methods

### Data compilation

Publications that reported data on litter mixture decomposition were selected from the Web of Science resource, Google Scholar, and the China National Knowledge Infrastructure. Keyword search strings consisted of term combinations, such as (litter or debris or residue) AND (mix* or diversity or non-additive effect or additive effect) AND (decompos* or decay* or degrad*). The PRISMA (Preferred Reporting Items for Systematic Reviews and Meta-Analyses) flow diagram performed the procedure used for the selection of studies for meta-analysis (Fig S2).

Each selected publication had to satisfy the following three criteria: 1) it had to report at least one of our selected variables [expected decay rate (Rexp) vs. observed decay rate (Robs), or species-specific decay rate in mixture Rmix vs. in unmixed (Rsin)], and the decay rate should been expressed as mass loss or mass remaining (%); 2) it had to provide the means and sample sizes (n) of the variables selected for the meta-analysis or could be calculated from the chosen papers; 3) the measurements of selected variables were performed at the same temporal and spatial scale; and 4) except litter mixture treatment, the study of other experimental treatments (such as nutrient addition, warming, water controlling) was excluded. If the data were presented in figures, Getdata Graph Digitizer (version 2.24) was used to extract the numerical values. In addition, we sent Emails to corresponding authors to query the original data of k values (yr^-1^) or relative mixture effect [(observed mass loss-expected mass loss)/ expected mass loss*100], when the selected variables used to estimate the non-additive effect were not provided in the papers.

We separated the dataset into two parts to perform two sub-meta-analyses (Fig. 1). The first sub-meta-analysis was to compare the decay rates of mixing litter between expected decay rates (Rexp) and observed decay rates (Robs). The second sub-meta-analysis was to compare the species-specific decay rates between single (Rsin) and mixture (Rmix) decomposition. As a note, the questions 1 to 5 were all answered by two sub-meta-analysis, the questions 6 and 7 were only answered by the second sub-meta-analysis. In total, 820 pair-observations in sub-meta-analysis 1 were obtained from 52 selected papers and 276 pair-observations in sub-meta-analysis 2 were obtained from 16 papers (Table S1). These studies were primarily conducted in East Asia, Western Europe, and North America (Fig. S3).

**Figure 1.**
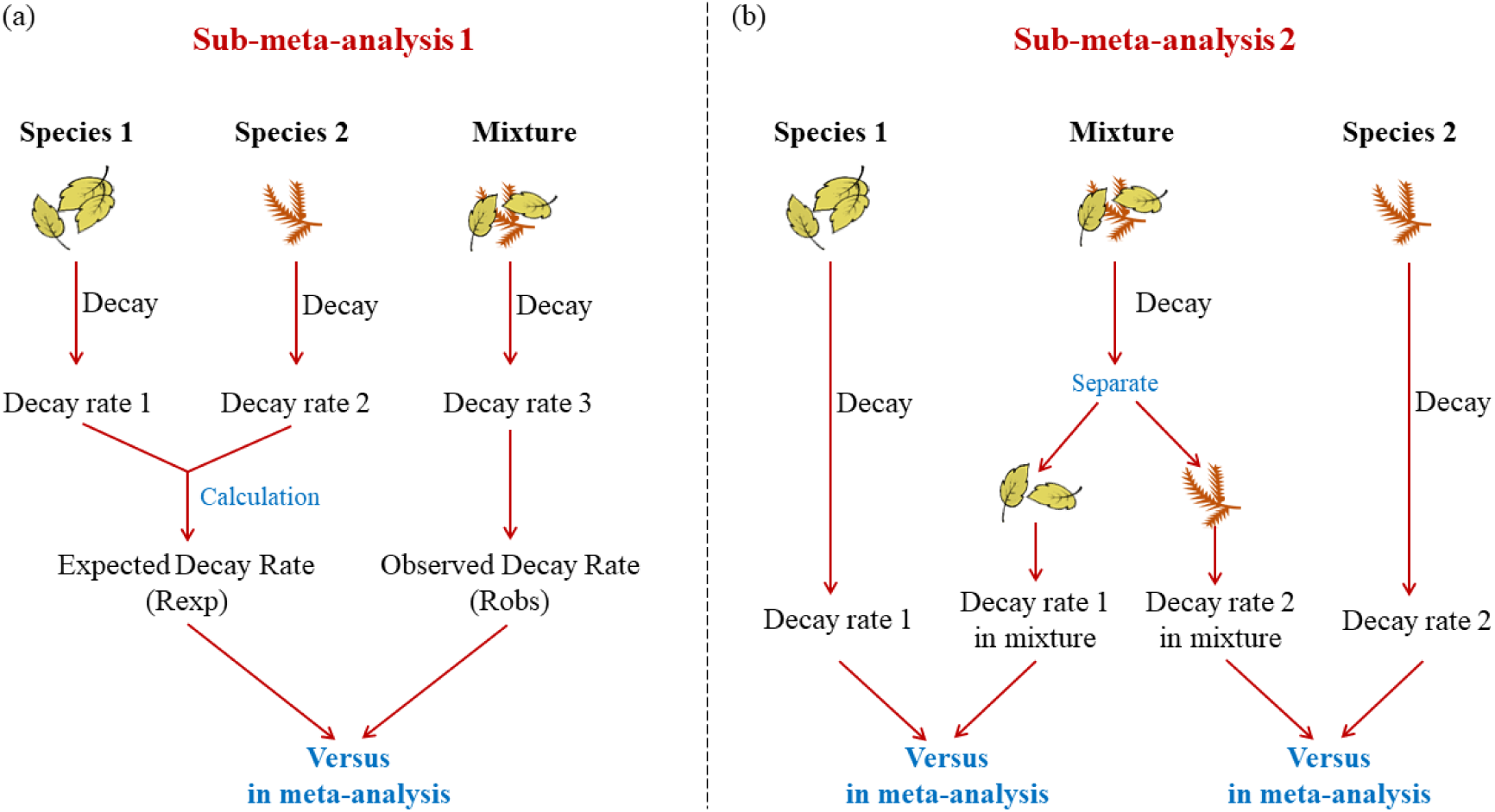
The two sub-meta-analyses in this study. (a) the sub-meta-analysis 1, comparison of mixing litter decay rate between expected rate (Rexp) and observed rate (Robs); (b) the sub-meta-analysis 2, comparison of species-specific decay rate between in single (Rsin) and in mixture (Rmix) decomposition. Rexp = w1R1+ w2R2+…+ wnRn, where wn is the weight of species n in the mixture and Rn is the decomposition rate of species n. The additive effect and non-additive effect are when “Rexp vs. Robs” or “Rsin vs. Rmix” are equal and not equal, respectively.

For each study, we also noted the factors relevant to the mixing effects including the site location, ecosystem type, species (and its nutrient contents if provided), decay period, mesh size, response variables, and other background information (e.g. mean annual temperatures (MAT), mean annual precipitation (MAP), etc.).

### Statistical analysis

The natural log-transformed response ratio (R), defined as the effect size (Hedges *et al.*, 1999), was used as an index to measure the effect level of litter mixture on decomposition rate:

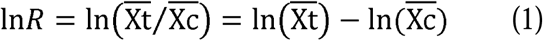

where 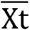 and 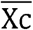 represent the Robs and Rexp, respectively; or the Rmix and Rsin, respectively.

The variance of ln*R* (*v*) was calculated using the following formula:

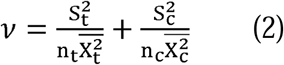

where S_t_ and S_c_ are the standard deviations (SDs) for the Robs and Rexp, respectively, or for the Rmix and Rsin, respectively; n_t_ and n_c_ are the sample sizes for the Robs and Rexp, respectively, or for the Rmix and Rsin, respectively. If both the SD and standard error (SE) were lacking, we estimated the missing SD by multiplying the average coefficient of variation (CV) from each data set by the reported mean value (Wiebe *et al.*, 2006).

A nonparametric weighting function was used to weight each individual study; and the mean effect size 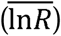 of all observations was estimated according to the Eq. 3:

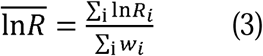

where *w* is the weighting factor used to calculate the inverse of the pooled variance (1/*v*); and lnR_i_ and w_i_ are the ln*R* and *w* of the *i*th observation, respectively.

To determine whether there was a significant difference under litter mixture treatment (Rosenberg *et al.*, 2000), we employed a fixed-effect model to calculate 95% confidence intervals (CIs) of the weighted effect size, using the Metawin 2.1 software. The effect was only considered significant if the 95% CI values did not overlap with 0. Furthermore, to clearly express the non-additive effects, the mean effect size was converted back to the percent change, using the following equation:

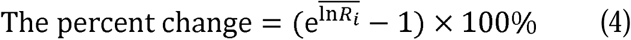

In order to better understand the formation mechanisms of the non-additive effects, we grouped the data according to climate zones (tropical, temperate, frigid), ecosystem types (forest, shrubland, grassland, aquatic, peatland), mesh sizes [small (diameter < 1 mm), medium (1mm≤diameter < 5) and large (diameter≥5mm)], and decay periods (< 180 days, 180-360 days, 360-720 days, and >720 days) (Knorr et al. 2005). Finally, we also grouped the trees and shrubs based on different functional types (broadleaf, needle, evergreen, and deciduous).

A continuous randomized-effect model was used to assess the potential linearity or non-linearity between ln*R* and climate factors (MAP and MAT)) or forcing factors (decay period, and initial nutrients content). The total ln*R*′ heterogeneity among the selected studies (Q_T_) was partitioned into different groups based on cumulative effect sizes (Q_M_) and the residual error (Q_E_) (Rosenberg *et al.*, 2000).

## Results

The mean effect size calculated across all the studies was significantly positive both in Rexp vs. Robs and Rsin vs. Rmix, with an average increase of +2% and +4%, respectively (Fig. 2a). When considering different climate zones, mixing litter decomposition caused a significantly positive response in temperate areas and a significant negative response in frigid areas in both of sub-meta-analyses (Fig. 2b). Unlike temperate and frigid areas, in tropical area the Rexp was significantly higher than Robs, and Rsin and Rmix differed little. The continuous randomized-effect model suggested that MAT had a significant positive correlation with the mean effect size in two sub-meta-analysises (Table 1). With respect to ecosystem types, the mean effect sizes of litter mixing on decomposition rate showed a significant positive response in Rexp vs. Robs for all five ecosystems except shrubland. (Fig. 2c).

**Table 1.** Relationships between the effect size of litter mixing on decay rate and mean air temperature (MAT), mean annual precipitation (MAP), experiment duration, species richness, and litter initial nutrients content.

| | $Q_T$ | $Q_M$ | $Q_E$ | Slope | *P*-value |
| --- | --- | --- | --- | --- | --- |
| **In sub-meta-analysis 1** |  |  |  |  |  |
| **MAT** | **575.37** | **28.88** | **546.49** | **0.004** | **<0.001** |
| MAP | 543.89 | 27.87 | 516.02 | <0.001 | <0.001 |
| Duration | 1192.93 | 2.63 | 1190.31 | <0.001 | <0.001 |
| Species richness | 1047.87 | 2.27 | 1045.60 | -0.008 | 0.13 |
| **In sub-meta-analysis 2** |  |  |  |  |  |
| **MAT** | **235.53** | **4.42** | **230.81** | **0.002** | **0.04** |
| MAP | 269.57 | 0.39 | 269.18 | <0.001 | 0.53 |
| Duration | 378.59 | 0.03 | 378.56 | <0.001 | 0.86 |
| Initial C | 285.88 | 2.99 | 282.89 | 0.003 | 0.08 |
| Initial N | 312.32 | 0.52 | 311.80 | -0.014 | 0.47 |
| **Initial P** | **137.69** | **3.86** | **133.83** | **-0.896** | **0.05** |
| Initial Lignin | 213.51 | 1.20 | 212.31 | 0.002 | 0.27 |
| Initial C/N | 282.12 | 0.14 | 281.98 | <0.001 | 0.71 |
| Initial C/P | 131.48 | 0.18 | 131.29 | <0.001 | 0.67 |
| Initial N/P | 132.68 | 0.19 | 132.49 | 0.001 | 0.66 |
| Initial N/L | 211.94 | 0.30 | 211.63 | -0.141 | 0.58 |
| Initial P/L | 51.77 | 0.42 | 51.34 | 14.735 | 0.52 |

Statistical results were reported as total heterogeneity in effect sizes among studies ( $Q_T$ ), the difference among group cumulative effect sizes ( $Q_M$ ), and the residual error ( $Q_E$ ) from continuous randomized-effects model of meta-analyses. The relationship is significant when $P < 0.05$ .

**Figure 2.**
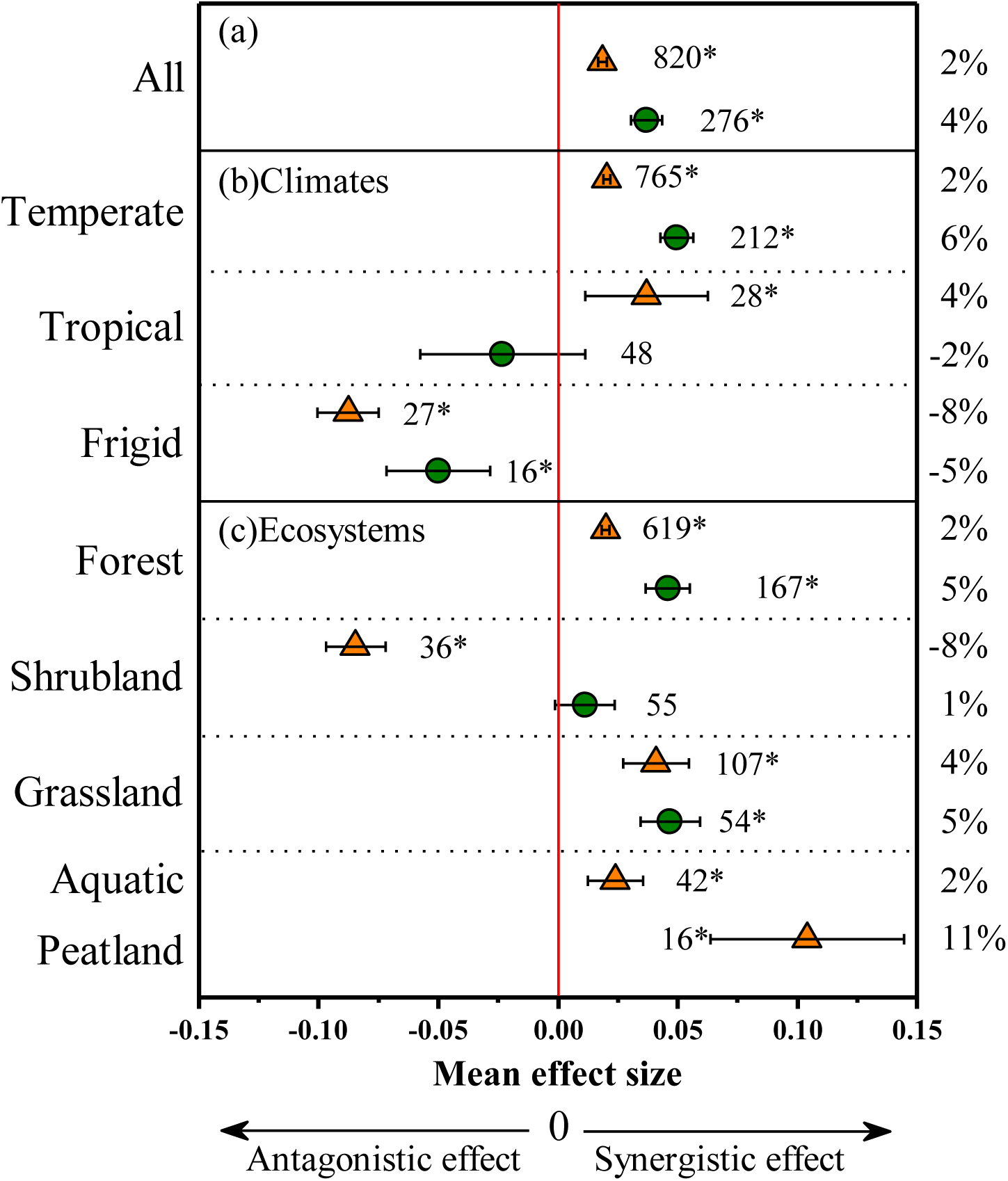
Comparison of mixing litter decay rates between expected values (Rexp) and observed values (Robs) (sub-meta-analysis 1, orange triangles), and comparison of species-specific decay rates between in single (Rsin) and in mixture (Rmix) decomposition (sub-meta-analysis 2, green circles) (a) across all studies, (b) among different climates, (c) ecosystems. If the mean effect size = 0, then means additive effect; If the mean effect size > 0, then means synergistic effect; If the mean effect size < 0, then means antagonistic effect. If the effect size 95% CI (error bars) did not cover zero, the non-additive effect was considered to be significant (*). The sample size for each variable is shown next to the point. The data on right-hand y-axis represent the mean percentage change for each variable (%).

The size level of the mesh in experimental methods resulted in different effects on the decomposition rate (Fig. 3). When mesh size was divided into three groups—small, middle, and large—and a marked antagonistic effect was found in studies of small mesh size as expressed by Rexp vs. Robs. In contrast, the decomposition rate in the studies of middle and large mesh sizes showed an obvious synergistic response (Fig. 3a). For Rsin vs. Rmix, the small and large mesh sizes depicted an additive effect, whereas, the middle mesh size showed a +6% increase in decomposition rate (Fig. 3a). When the decay period was partitioned into four levels, results showed that the decomposition rate increased under short-term duration (<180 days) and medium-term (180-360 days and 360-720 days), instead, a significant antagonistic effect was observed under long-term duration (>720 days; -4%) in Rexp vs. Robs (Fig. 3b).

**Figure 3.**
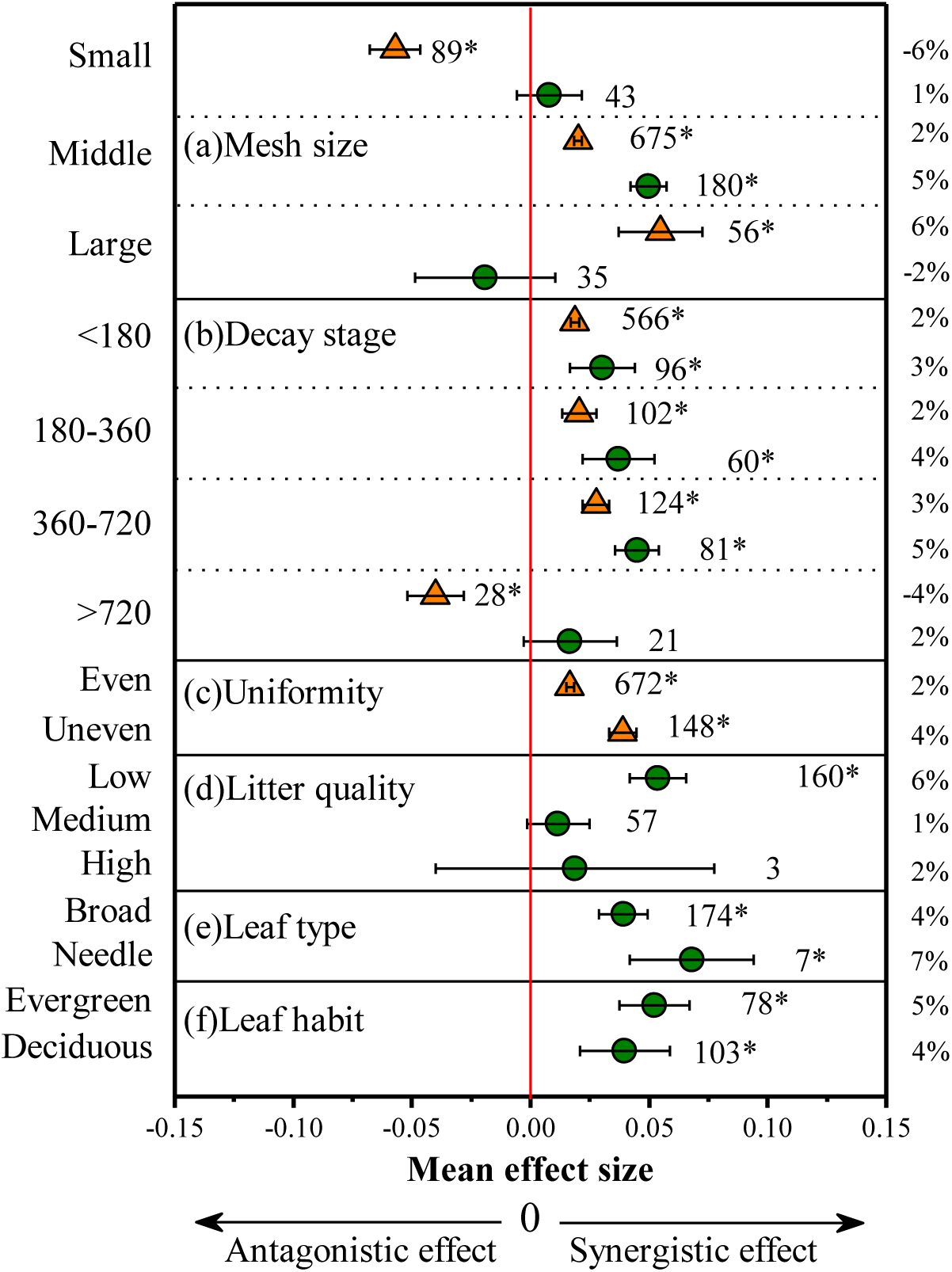
Comparison of mixing litter decay rates between expected values (Rexp) and observed values (Robs) (sub-meta-analysis 1, orange triangles), and comparison of species-specific decay rates between in single (Rsin) and in mixture (Rmix) decomposition (sub-meta-analysis 2, green circles) among different experimental methods. (a) mesh size, (b) duration, (c)uniformity, (d) litter quality, (e) leaf types, and (f) leaf habit. If the mean effect size > 0, then means synergistic effect; If the mean effect size < 0, then means antagonistic effect. If the effect size 95% CI (error bars) did not cover zero, the non-additive effect was considered to be significant (*). The sample size for each variable is shown next to the point. The data on right-hand y-axis represent the mean percentage change for each variable (%).

In sub-meta-analysis 1 (Rexp vs. Robs), both of even litter mixtures and uneven litter mixtures showed synergistic effects on decomposition rate (Fig. 3c). In sub-meta-analysis 2 (Rsin vs. Rmix), the litter quality was divided into three levels based on lignin content: low, medium, and high—and the results showed that the decomposition rate of low-quality litter exhibits a greater positive change than the middle and the high litter (Fig. 3d). When tree species were grouped into broad/needle or evergreen/deciduous, results consistently showed synergistic effects. Besides, the continuous randomized-effect model of meta-analyses showed a significant positive correlation between litter initial P and the effect size of the sub-meta-analysis 2 (Rsin vs. Rmix), yet, the remaining initial nutrients of litter did not show any correlation (Table 1).

## Discussion

This study is primarily dedicated to calculate the general effects of litter mixture on its decay rates. Our results showed that the litter mixture widely demonstrates a non-additive effect, and most frequently synergistic effect, which is consistent with two previous review studies (Gartner & Cardon 2004; Li *et al.*, 2016). Specifically, the decay rates of litter mixture are on average 2%-4% faster when compared to that of single litter species (Fig. 2a). This significant synergy is weak but logical. When different litter species mixing, many processes (including the stimulating and restraining) occur simultaneously with one counterbalancing the others more or less. Based on our meta-analysis results, we further raised a conceptual model for the non-additive effect of mixed litter samples of two species (Fig. 4). It should be noted that the model reflects a general pattern, which may not be appropriated in all cases.

**Figure 4.**
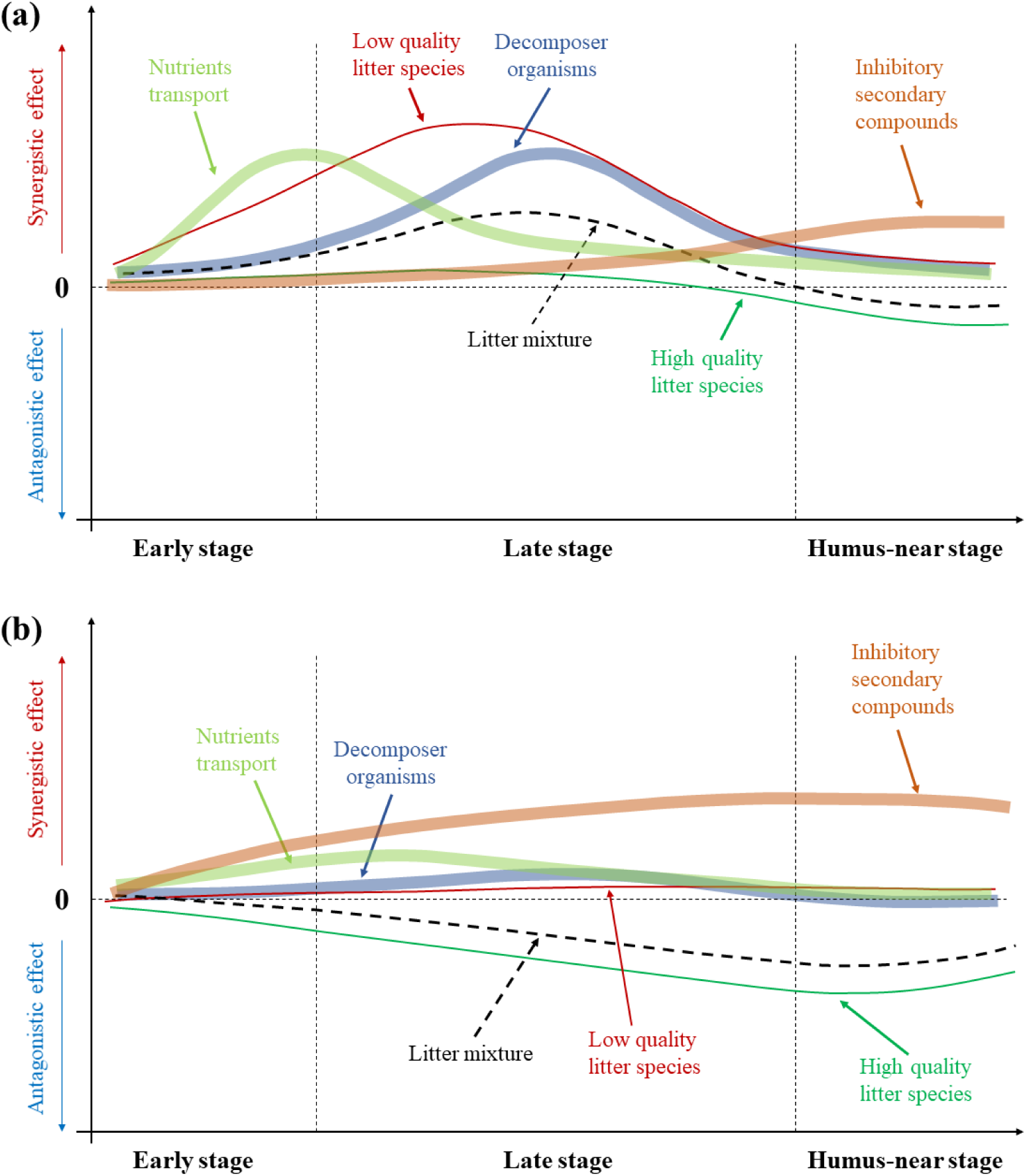
Conceptual model for non-additive effect in mixed litter decomposition of two species based on our meta-analysis. The three-stages referred to (Berg 2014). (a) The non-addition effect in temperate and tropical areas; (b) The non-addition effect in frigid areas or absence of soil fauna.

The variations of litter decomposers (including microorganisms and soil fauna) play an important role in non-additive effect. Although our meta-analysis lacked sufficient data to discuss the decomposer variations, the finding (mean effect size = 0.1604; 95% CI: 0.1432∼0.1776, Table S2) suggested that microbial biomass is significantly higher in litter mixture compared to single litter. In the decomposition of litter mixture, on one hand, high-quality litter brings more available carbon to support microbes growth (Hättenschwiler & Jørgensen 2010), on the other hand, microbial activities and extracellular enzyme activities also increase simultaneously (Hu *et al*. 2006; Cong *et al*. 2015; Gogo *et al*. 2016), both of which can help the decay of low-quality litter to generate synergistic effect.

When the data were divided into different climatic zones, an antagonistic effect of litter mixture was observed in frigid areas in the two sub-meta-analysis (Fig.2b and 4b, answer to Q1). Furthermore, a significant positive correlation between MAT and its effect size on decomposition rate also supported this hypothesis (Table 1). This phenomenon likely resulted from 1) soil fauna and microbial biomass being low in high latitude regions compared with low latitude regions (Fierer *et al.*, 2009; Xu *et al.* 2013; Nielsen *et al.* 2014); 2) the soil organisms’ activities being limited by lower temperatures in high latitude areas. As mentioned before, soil fauna are regarded as an important propelling force for the synergistic effect, once their quantity and activities are substantially reduced, a synergistic effect seems unable to develop; and 3) the fungi hyphae growth being limited by low temperature and less rain in frigid area impeding nutrients transport from high-quality litter to the low-quality. When the data were classified into different ecosystems, only the shrubland showed an antagonistic effect (Fig.2c, answer to Q2), which would be because that most of the shrubland data were acquired from frigid areas.

Soil fauna is an important group of decomposers but difficult to directly control in the studies of litter decomposition. The size of mesh diameter is usually introduced to distinguish different kinds of soil fauna. While a fine mesh (<1mm) is applied to exclude most of the soil fauna, the antagonistic effect plays a leading role (Fig. 3a and 4b, answer to Q3). Interestingly, the antagonistic effect will turn into a synergistic effect when middle (1-5 mm) or large mesh (>5 mm) is used. This emerging conclusion was also drawn in a recent litter mixing study (Barbe *et al*. 2018). High-quality litter with rich nutrients and energy would be palatable to soil fauna (Zhang *et al*. 2016), which may accelerate the decay rate of litter mixture. A global synthesis studies also suggested that the decay rate of litter fall by a third if the soil fauna excluded (Zhang *et al*. 2015). Therefore, the existence of high-quality litter in mixing litter decomposition can promote litter decay rate. In addition, the soil fauna may make the litter more accessible to bacteria and fungi, which could stimulate the microbial growth and therefore decomposition further (Smith & Bradford 2003). An inexplicable phenomenon for the present is that a non-significant antagonistic effect occurred when large mesh was used in sub-meta-analysis 2, which could be brought by the error of insufficient data (35 observations). In order to determine whether these results develop consistently, more extensive studies are required.

It is worth noting that the synergistic effect gradually weakens as a function of decomposition time and turns into an antagonistic effect after 720 days of decay (Fig. 3b and 4a, answer to Q4). In the early stage of decomposition, the input of fresh litter provides abundant food for the soil fauna and microorganisms. In addition, studies suggest that nutrients transport frequently occurs in the early stage of decomposition (Hansson *et al*., 2010; Liao *et al*., 2016). Thus, the rich food combined with high-efficient nutrient transport may facilitate the growth and activities of soil microorganisms, which ultimately produces a synergistic effect. As stated before, the effects of soil fauna are important to the synergistic effect, however, their relative role maybe be weaken and their effects maybe disappear at late stage of decomposition. Consequently, the litter decomposition is mainly performed by the microbes that able to degrade very recalcitrant compounds (such as lignin, tannin, etc.) (Guénon *et al*. 2017, Chapman *et al.* 2013). Besides, the remaining recalcitrant substances from low-quality litter may resist the degradation at late stage of decay and subsequently generate an antagonistic effect (Fig. 4).

Both of species evenness and richness influence the ecosystem functions, including litter decomposition processes (Dangles & Malmqvist 2004; Tilman *et al*., 2014). Although the species evenness depicts inconsistent effects on litter decomposition (Hillebrand et al., 2008; Ward *et al*., 2010; Li *et al*., 2013), this meta-analysis indicates that the synergistic effect of uneven litter mixtures is higher than that of even mixtures (Fig. 3c, answer to Q5). We speculated that natural field values of uniformity reflected in uneven mixtures more favorable for the microorganisms’ growth than in even mixtures (Swan *et al*. 2009). This opinion requires further investigations to be confirmed. Our results also show a negative but insignificant relationship between litter richness and its effect size on litter decomposition in sub-meta-analysis 1 (Table 1, answer to Q5). Studies suggested that fungal diversity increased with litter richness (Otsing *et al*. 2018), but we still do not know how fungi change influences the decomposition of litter mixture. Although the underlying reasons are unclear, we speculate that the following two points may contribute to this phenomenon. First, the rate of litter decomposition might have a maximum value (Fig. 4) (Barbe *et al*. 2018), the non-additive effect would not be permitted to beyond this maximal decomposition with litter richness. Second, the small size of data is not enough to reveal the true rule.

With respect to the three classifications of litter species in sub-meta-analysis 2, a noteworthy phenomenon is that a significant synergistic effect was observed in low-quality litter species, while no significant change detected in medium and high-quality litter species (Fig 3d and 4a, answer to Q6). Although the underlying reasons are unclear, we assumed that the following three factors may contribute to this result. First, the nutrients released from high-quality litter species promote low-quality litter decay (Versini *et al*. 2016). Second, the input of high-quality litter species promotes the growth of soil organisms (Hättenschwiler & Jørgensen 2010, Chapman *et al*. 2013), thus leading to low-quality litter becoming competitive and accelerating its decomposition. Third, the improved microenvironment through the addition of high-quality litter, facilitates the decomposition of low-quality litter (He *et al.*, 2019). High initial N and P content of litter indicate high decomposability. The strong negative relationship between the mean effect size and litter initial N (*P* > 0.05) and P (*P* < 0.05) also support above result (Table 1). But we need more data to further clear and definite this relation.

As the trees and shrubs were classified into four groups: broad, needle, evergreen, and deciduous, the results showed that the mean effect size of needle and evergreen groups were higher than those of broad and deciduous groups (Fig 3e, f, answer to Q7). In general, the needle species contain more lignin than broadleaf species, and they will act as a beneficiary of synergistic effect in litter mixing. Mean lignin content of evergreen leaves was 26% in this sub-meta-analysis database, which is higher than that of deciduous leaves (23%, Table S1). Moreover, the deciduous leaves contain more N and P than evergreen leaves based on the large scale research (Han *et al*. 2005), meaning the decomposability of deciduous leaves is greater than that of the evergreen leaf (Cornwell *et al.*, 2008; Liu *et al.*, 2016). The data indicates that the decomposition of low-quality litter species is more sensitive to the treatment of litter mixing, which is in agreement with Q6.

## Conclusions

In summary, litter mixing generally increased the decay rate and tended to cause a synergistic effect compared to single litter. Moreover, when the mixing litter was separated, low quality-litter species displayed a synergistic effect, yet there was no change in high-quality litter species. A synergistic effect usually occurred at the early and late decay stage and disappeared at the humus-near stage. The soil organisms, especially soil fauna, were regarded as the important factors generating a synergistic effect. We suggest that, the synergistic effect and antagonistic effect, whose interplay give rise to the non-additive effect, occur simultaneously rather than independently. Whereas, some of our results have not been confirmed so far at the scale of a meta-analysis. Additional investigations are still needed to improve both the theory and the model in the near future.

## Acknowledgements

We thank all the authors of the original studies included in our meta-analysis. The study was supported by the National Natural Science Foundation of China (41703065, U1805243, 41807028, 31460077), the Natural Science Foundation of Fujian Science and Technology Department (2018J01621), and the Education and Research Projects for Young Teachers of Fujian Provincial Education Department (JAT170187).

## Author contribution

J. L, Y. H and Q. Y. designed the study. J. L conducted data checking. J. L and Q. Y. conducted the analyses. J. L and Y. H wrote the first draft of the manuscript. J. L., H. W., X. L., Q. S., F. L., Y. H., Q. Y. contributed to data collection, writing, and revision.

## Supporting Information

**Figure S1** The non-additive effect in mixing litter decomposition.

**Figure S2** PRISMA diagram showing the process of locating studies include in this meta-analysis

**Figure S3** World distribution of selected studies in this meta-analysis.

**Note S1** List of all the references used in the meta-analysis.

**Table S1** Summary of the references and data used in the meta-analysis of the effects of litter mixing on decomposition rate.

**Table S2** Summary of the data and result used in meta-analysis of the effect of litter mixing on microbial biomass.

