## Supplementary material for "Synergistic effect: a common theme in mixed-species litter decomposition": Fig. S1-S3

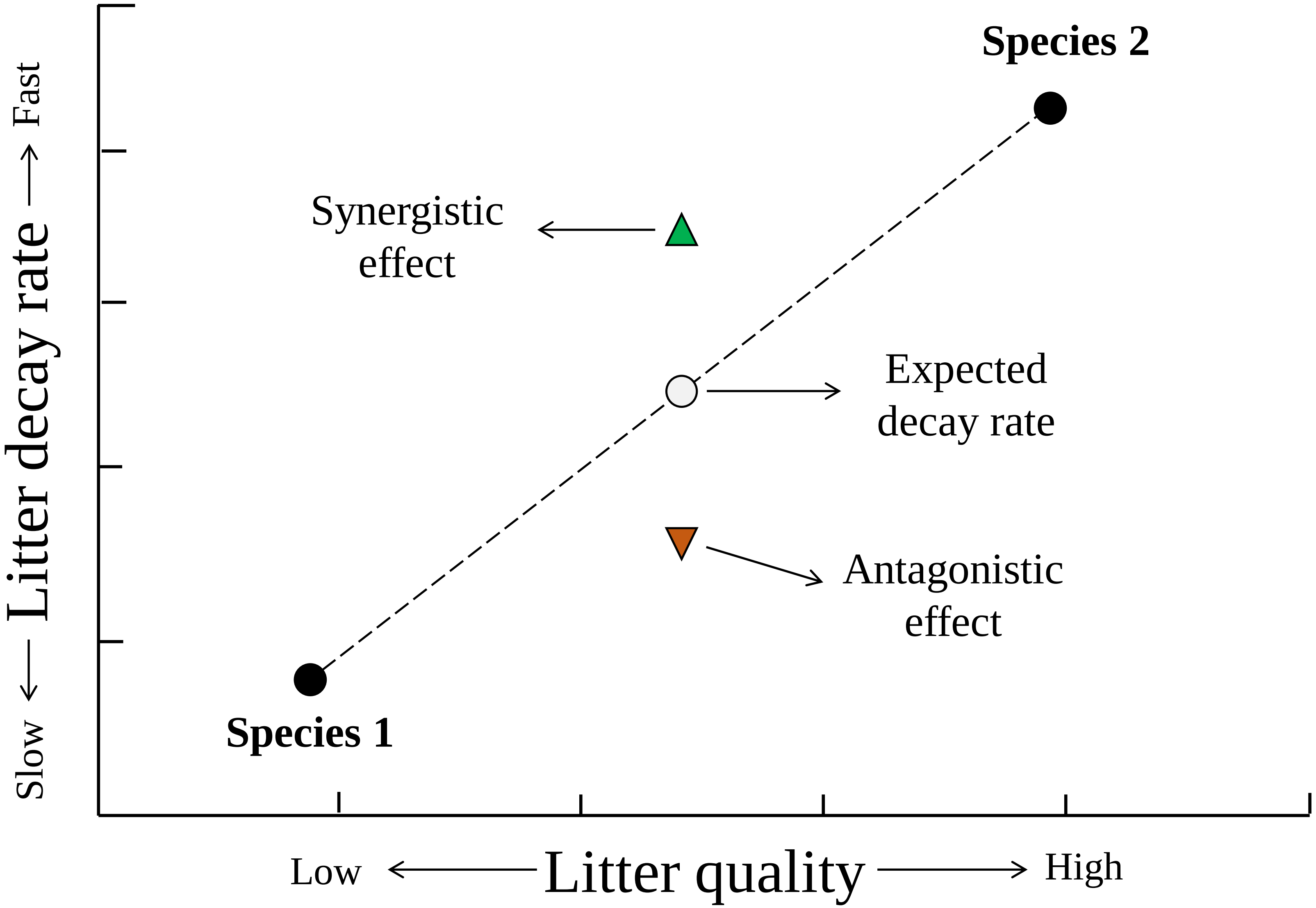


**Figure S1** The non-additive effect in mixing litter decomposition. The non-additive effect includes the synergistic effect (green triangle) and antagonistic effect (yellow triangle).

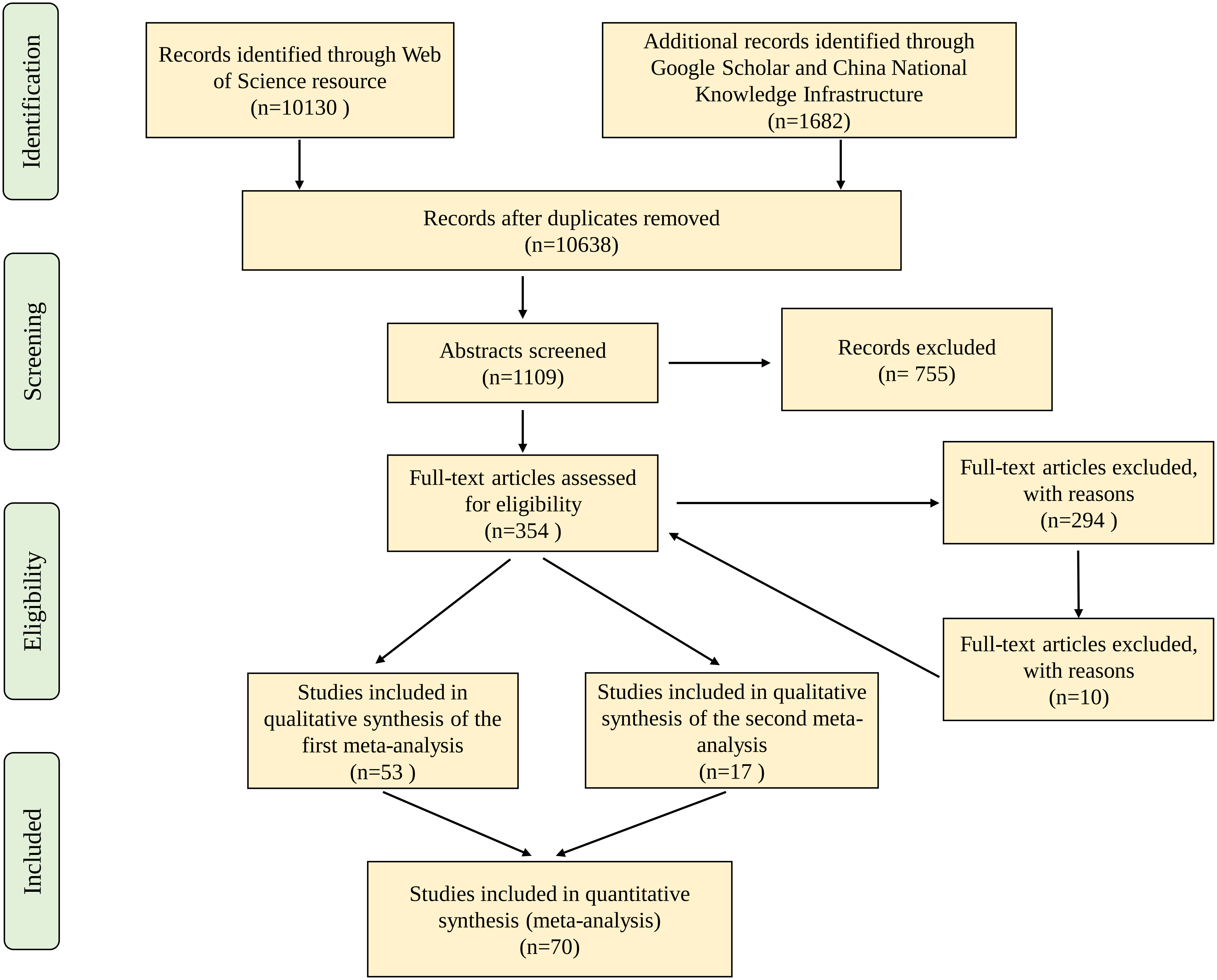


Figure 2s PRISMA diagram showing the process of locating studies include in this meta-analysis

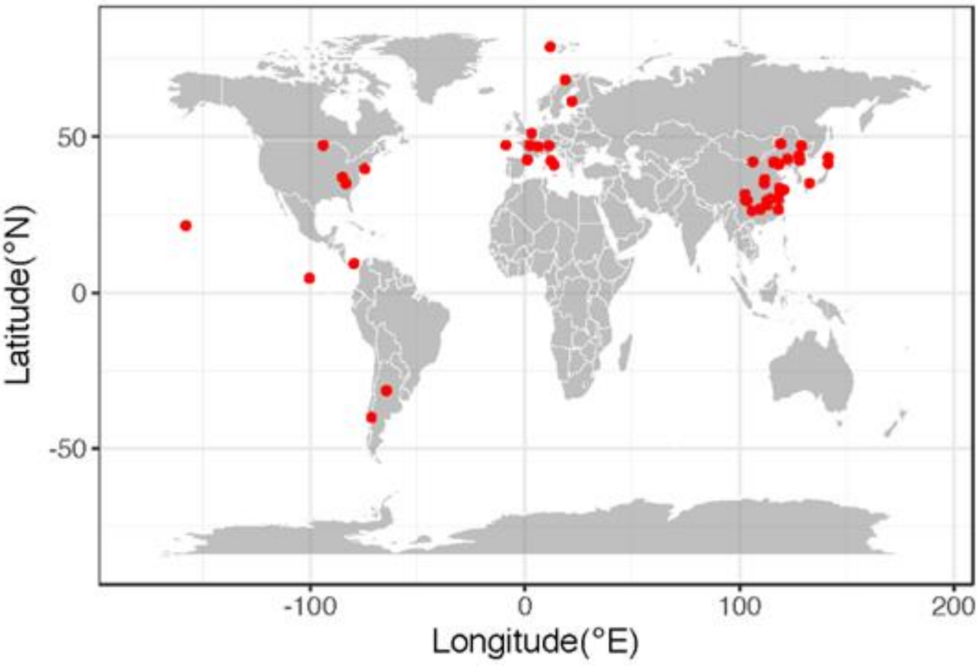


**Figure S3** World distribution of selected studies in this meta-analysis.
